# Subcortical Hubs of Brain Networks Sustaining Human Consciousness

**DOI:** 10.1101/2024.10.04.616722

**Authors:** Morgan K. Cambareri, Andreas Horn, Laura D. Lewis, Jian Li, Brian L. Edlow

## Abstract

Neuromodulation of subcortical network hubs by pharmacologic, electrical, or ultrasonic stimulation is a promising therapeutic strategy for patients with disorders of consciousness (DoC). However, optimal subcortical targets for therapeutic stimulation are not well established. Here, we leveraged 7 Tesla resting-state functional MRI (rs-fMRI) data from 168 healthy subjects from the Human Connectome Project to map the subcortical connectivity of six canonical cortical networks that modulate higher-order cognition and function: the default mode, executive control, salience, dorsal attention, visual, and somatomotor networks. Based on spatiotemporally overlapped networks generated by the Nadam-Accelerated SCAlable and Robust (NASCAR) tensor decomposition method, our goal was to identify subcortical hubs that are functionally connected to multiple cortical networks. We found that the ventral tegmental area (VTA) in the midbrain and the central lateral (CL) and parafascicular (Pf) nuclei of the thalamus – regions that have historically been targeted by neuromodulatory therapies to restore consciousness – are subcortical hubs widely connected to multiple cortical networks. Further, we identified a subcortical hub in the pontomesencephalic tegmentum that overlapped with multiple reticular and extrareticular arousal nuclei and that encompassed a well-established “hot spot” for coma-causing brainstem lesions. Multiple hubs within the brainstem arousal nuclei and thalamic intralaminar nuclei were functionally connected to both the DMN and SN, emphasizing the importance of these cortical networks in integrative subcortico-cortical signaling. Additional subcortical connectivity hubs were observed within the caudate head, putamen, amygdala, hippocampus, and bed nucleus of the stria terminalis, regions classically associated with modulation of cognition, behavior, and sensorimotor function. Collectively, these results suggest that multiple subcortical hubs in the brainstem tegmentum, thalamus, basal ganglia, and medial temporal lobe modulate cortical function in the human brain. Our findings strengthen the evidence for targeting subcortical hubs in the ventral tegmental area, central lateral thalamus, and pontomesencephalic tegmentum to restore consciousness in patients with DoC. We release all subcortical connectivity maps to support ongoing efforts at therapeutic neuromodulation of consciousness.

## Introduction

Human consciousness requires functional connections between subcortical and cortical networks that mediate arousal and awareness, respectively (Edlow et al., 2024; Koch et al., 2016). In patients with severe brain injuries, the reintegration of connectivity between subcortical and cortical networks is essential for recovery of consciousness (Edlow, Claassen, et al., 2021). However, there are currently few therapies proven to promote recovery of consciousness in patients with severe brain injuries (Giacino et al., 2012), a limitation in clinical care that is partly attributable to a lack of therapeutic targets (Edlow, Sanz, et al., 2021). To identify such targets, it is essential to generate a reliable map of subcortical-cortical connectivity in the healthy, conscious human brain, as this map may be used to guide the search for widely connected network hubs that could be stimulated to restore consciousness in the injured brain.

Growing evidence suggests that widely connected network hubs play a key role in higher-level cognitive functions (van den Heuvel & Sporns, 2013) and that disconnection of network hubs is implicated in the pathogenesis of a broad range of neuropsychiatric disorders (Crossley et al., 2014). In patients with disorders of consciousness (DoC), the relevance of network hubs to the loss and restoration of consciousness is supported by human studies showing that hub lesions cause coma (Edlow et al., 2013; Fischer et al., 2016; Parvizi & Damasio, 2003) and that hub stimulation may promote the reemergence of consciousness (Giacino et al., 2012; Schiff et al., 2007). Animal models have similarly revealed that focal lesions within the brainstem tegmentum (Fuller et al., 2011; Pais-Roldan et al., 2019) and targeted stimulation within the central thalamus (Redinbaugh et al., 2020; Tasserie et al., 2022) may cause loss and restoration of consciousness, respectively, under a variety of experimental conditions. Consistent with these animal models, clinical trials in humans with DoC have historically targeted subcortical hubs in the central thalamus (Cain, Spivak, et al., 2021; Cain et al., 2022; Schiff et al., 2007), brainstem tegmentum (Edlow et al., 2020; Elias et al., 2021; Fridman et al., 2019; Giacino et al., 2012), and basal ganglia (Cain, Visagan, et al., 2021; Whyte et al., 2014).

A barrier to progress in therapeutic target selection for humans is that there are gaps in knowledge about how subcortical hubs modulate their cortical counterparts in the healthy, conscious human brain. While extensive progress has been made in understanding this physiology in animals (Aston-Jones et al., 2001; Moruzzi & Magoun, 1949; Scammell et al., 2017; Steriade & Glenn, 1982; Vertes & Martin, 1988) via electrophysiological, lesion-based and tract-tracing studies, the physiology of subcortical networks has not been fully elucidated in humans. Specifically, it is unknown which subcortical hubs activate cortical networks, inhibit cortical networks, or mediate state switches. Identifying integrative network hubs that link arousal and awareness may reveal the subcortical targets most likely to promote recovery of consciousness in patients with DOC.

Advances in functional MRI (fMRI) (Luppi et al., 2024) and the availability of large normative datasets as part of the Human Connectome Project (HCP) (Glasser et al., 2016; Smith et al., 2013) now create opportunities to identify subcortical network hubs and elucidate the mechanisms by which they modulate human consciousness, providing new therapeutic target candidates for patients with DoC. Here, we aimed to identify the subcortical regions of the human brain that possess high levels of functional connectivity with multiple cortical networks – regions that we define as subcortical network hubs. While this approach to functional connectivity mapping cannot determine the direction of electrical signaling between subcortical and cortical regions, there is evidence that stimulating small subcortical regions is an effective strategy for activating multiple cortical networks via diffuse ascending connections (Horn & Fox, 2020).

Using the NASCAR decomposition method on the resting-state fMRI (rs-fMRI) data from 168 HCP healthy subjects, we generated subcortical functional connectivity maps for six canonical large-scale brain networks. We identified subcortical hubs that strongly connect to multiple cortical networks by measuring the overlaps in functional connectivity maps within the subcortical regions. Further, we characterized the functional connectivity properties by which subcortical hubs interact with cortical networks (i.e., correlated versus anticorrelated functional connectivity). We release all functional connectivity maps to support progress in the field of human brain mapping and to inform the design of future clinical trials aimed at neuromodulation of consciousness.

## Materials and methods

### 7 Tesla rs-fMRI data

We analyzed the minimally preprocessed 7 Tesla (7T) rs-fMRI healthy control data (Glasser et al., 2013) from the WashU/U Minn component of the HCP (Glasser et al., 2016). Of the 176 subjects in the dataset, 168 were included due to reported acquisition and preprocessing issues in 8 subjects. Each rs-fMRI dataset was collected in four independent 15-minute sessions using a gradient-echo EPI sequence (1.6 mm isotropic voxels, TE = 22.2 ms, TR = 1000 ms) with opposite phase encoding directions (AP, PA). We used the first session with PA phase encoding direction to minimize inter-subject misalignment, consistent with recent work (Li, Curley, et al., 2021). These data were coregistered to the MNI 152 6th-generation space and represented in CIFTI grayordinate space with 91,282 total cortical and subcortical vertices/voxels (approximately 32K vertices per hemisphere cortically and an additional 32K voxels in the subcortex) as part of the HCP minimal preprocessing pipeline (Glasser et al., 2013). No additional smoothing was applied to avoid blurring across different functional regions (Li et al., 2018).

### Identification of spatially overlapped brain networks using NASCAR

We used the Nadam-Accelerated SCAlable and Robust (NASCAR) tensor decomposition method to identify large-scale brain networks. The key distinction between NASCAR and traditional seed-based methods (Lee et al., 2013) or parcellation schemes (Yeo et al., 2011) is that NASCAR allows networks to be spatially overlapped and temporally correlated. Unlike other commonly used data-driven approaches, such as the independent component analysis (ICA) or principal component analysis (PCA), NASCAR does not impose either independence or orthogonality constraints. Recent evidence suggests that networks identified by NASCAR are more physiologically plausible than ICA or PCA (Li et al., 2023; Li, Wisnowski, et al., 2021) and that subcortical functional connectivity identified by NASCAR closely corresponds to gold-standard histopathology (Li, Curley, et al., 2021). The allowance of spatial overlaps between networks enables the identification of subcortical network hubs, which are defined as regions that participate in multiple cortical networks.

#### Temporal synchronization using group BrainSync

Prior to NASCAR, an inter-subject temporal synchronization method, “BrainSync” (Anand et al., 2018), was performed to align the rs-fMRI signals across subjects. This step was necessary as rs-fMRI data reflect spontaneous brain activity and are not synchronous between subjects. To account for potential bias in the synchronization target, we used the group version of the BrainSync algorithm (Akrami et al., 2019), creating a virtual reference subject, which is, on average, close to all subjects in the population. Each subject’s data was then temporally aligned to that virtual reference subject. Critically, this synchronization procedure does not change the subject connectivity profile, as it would be measured by spatial correlations. The group result can then be mapped back to the individual space, if desired, using the invertibility property of the BrainSync algorithm.

#### NASCAR tensor decomposition

The synchronized rs-fMRI data were concatenated along the subject dimension, forming a third-order tensor, which has a spatial dimension (number of total vertices/voxels) of 𝑉 = 91292, a temporal dimension (number of time points) of 𝑇 = 900, and a subject dimension (number of subjects) of 𝑁 = 168. We ran NASCAR on this third-order tensor and obtained low-rank components representing large-scale brain networks common across all subjects.

### Identification and classification of functional brain network hubs

#### Identification of whole-brain resting state networks from the cortical maps

Each spatial map identified using the NASCAR method represents the spatial distribution of a whole-brain network consisting of both the cortical component and the subcortical component. Fig. 1 shows an example network encoded in the CIFTI format. The network map was separated into two cortical (left and right hemispheres) and one subcortical component and mapped onto the surface and the volumetric space for visualization, respectively. We visually examined the cortical components and identified six canonical resting-state networks, all consistent with prior literature (Raichle, 2011; Yeo et al., 2011), including the default mode network (DMN), the salience network (SN), the executive control network (ECN), the dorsal attention network (DAN), the somatomotor network (SMN), and the visual network (VN) (Fig. S1). Although a potential limbic network with medial temporal and temporal pole signals was visually identified, this component did not have connectivity within the orbitofrontal cortex, as previously reported in (Yeo et al., 2011). Additionally, a recent fMRI study suggested that the limbic network is part of the extended DMN rather than a separate and functionally distinct large-scale network (Girn et al., 2024). Therefore, we focused on the abovementioned six major large-scale networks and did not include a separate limbic network for the following analyses

**Figure 1:**
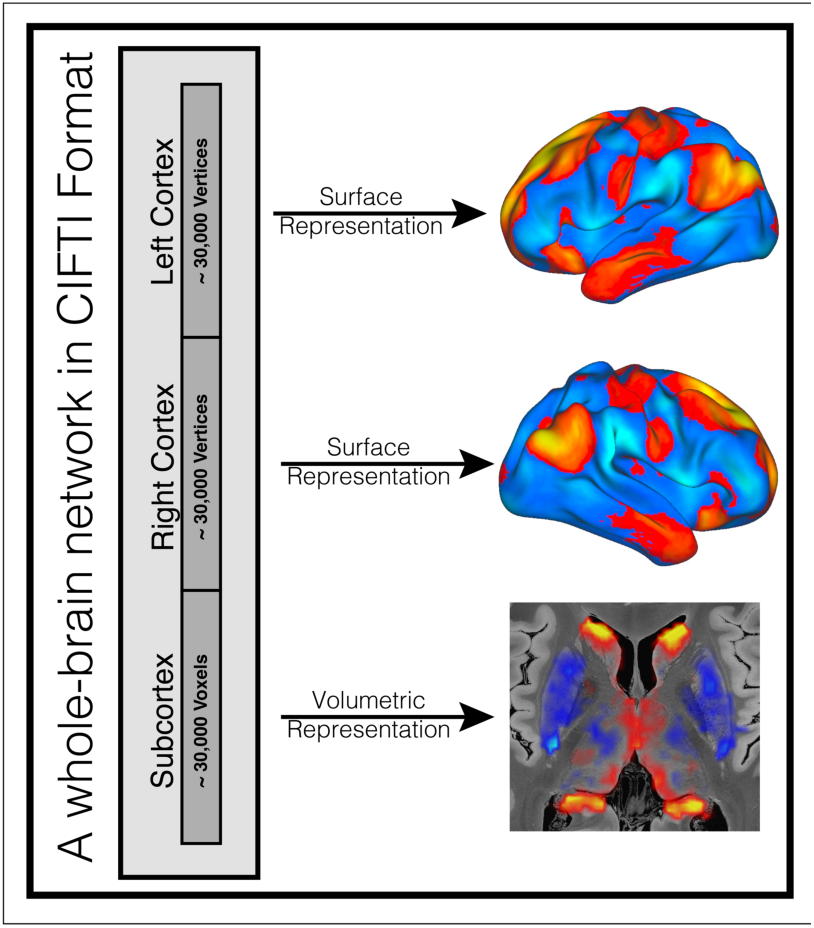
An example network (DMN) identified by the NASCAR method. The spatial map represented in the CIFTI format consists of both cortical and subcortical components. They were separated and mapped onto the surface and the volumetric space for visualization, respectively.

#### Separation of subcortical resting state networks

For each network, we extracted the subcortical component (Fig. 1 bottom) and converted them to a 3D volume. We then transformed this volume into the MNI NLIN 2009 space and linearly interpolated the map to 0.5 mm^3^ resolution. We removed the cerebellar signal from the subcortical volumes for this study because the cerebellum is closer to the head coil and has a higher signal-to-noise ratio (Koike et al., 2021) compared to the subcortical structures we are focusing on in this work, thus could introduce a strong bias towards the thresholding procedure below. Furthermore, while damage to the brainstem, thalamus, and basal ganglia have been associated with DoC, individuals with damage to the cerebellum (or born without a cerebellum) can sustain arousal and awareness (Lemon & Edgley, 2010), which makes the cerebellum less therapeutically relevant in patients with DoC.

#### Identification of subcortical resting state network hubs

Our first goal was to identify subcortical nodes whose BOLD signal was correlated or anticorrelated with multiple cortical networks. We identified the “correlated network hubs” as follows: We first binarized the subcortical map for each individual network by thresholding their distribution to preserve only the top 5% of values and then superimposed the six binarized maps. We identified the “anticorrelated network hubs” similarly using the bottom 5% threshold. Mixed hubs were identified by combining the results from the correlated and anticorrelated network hub overlaps. See Table 1 for the full descriptions of each hub type.

**Table 1:** Definition of three types of network hubs.

| **Classification** | **Description** |
| --- | --- |
| “Correlated” network hubs | Each network map was thresholded to preserve the top 5% of its most correlated voxels within the subcortex (i.e., the top 5% of the distribution), creating a binary network mask. Each correlated network mask was overlapped with one another to identify cortical regions that are highly correlated to multiple cortical networks. |
| “Anticorrelated” network hubs | Each network map was thresholded to preserve the top 5% of its most anticorrelated voxels within the subcortex (i.e. the bottom 5% of the distribution), creating a binary network mask. Each anticorrelated network mask was overlapped with one another to identify subcortical regions that are highly anticorrelated to multiple cortical networks. |
| “Mixed” network hubs | Each network map was thresholded to preserve the top 5% of its most correlated and anticorrelated voxels within the subcortex (i.e., both the top 5% and bottom 5% of the distribution), creating a single mixed binary network mask. Each mixed network mask was overlapped with one another to identify subcortical regions that have strong relationships to multiple cortical networks but with mixed signs. |

In addition to visualizing the neuroanatomic overlap of binarized network maps, we measured the “hubness” at each voxel by summing how many networks “passed” (above for correlated hubs and below for anticorrelated hubs) the 5% threshold (i.e., accumulating the binary masks across networks). This created a discrete hub map with values ranging from 0 to 6 (“0” means no network survived the 5% thresholding at this location, and “6” means all networks survived). We repeated this hubness measure for anticorrelated network maps. Furthermore, we identified subcortical nodes whose BOLD signal was highly connected to multiple cortical networks regardless of sign (correlated or

anticorrelated). To do so, we measured the hubness using the binarized “mixed hub” maps and summed the number of networks falling into either the top 5% or the bottom 5% of the distribution at each voxel.

#### Network threshold contours

To have a more granular and contiguous view of the network hubs, we approached the same question from a different perspective. For a fixed 𝑘 number of networks at each voxel, we queried the highest threshold that could be used to binarize the network maps such that there are 𝑘 networks overlapped at this location. We refer to these continuously valued threshold maps as “contour maps” hereafter. Note that in this analysis, different network combinations could occur at different locations. We use 𝑘 = 4 for visualization trade-off to best illustrate the consistency and correspondence between our hubness results and regions historically associated with consciousness level. All images in this paper are shown in radiological convention where the left hemisphere is shown on the right side of the image and vice versa.

## Results

### Visualization of subcortical resting state networks

Fig. 2 shows the subcortical maps of six canonical resting-state networks identified by the NASCAR decomposition method at axial slices through the pons, midbrain, diencephalon, basal forebrain, and basal ganglia. These network maps are available to view at https://github.com/ComaRecoveryLab/Subcortical_Network_Hubs. Visual inspection revealed that the networks were mostly symmetric across the midline of the brain, with occasional hemispheric differences in the connectivity profile.

**Figure 2:**
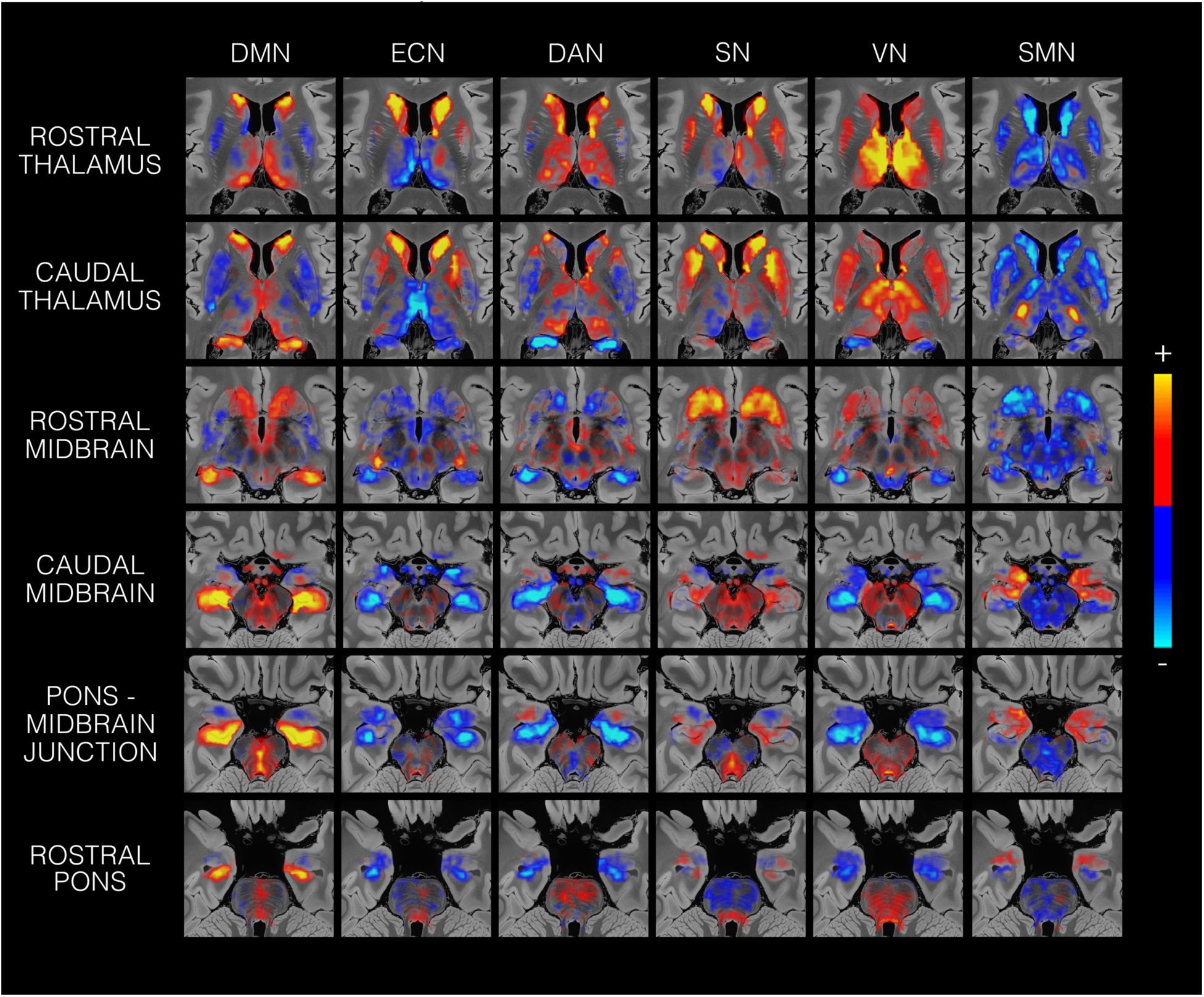
Subcortical connectivity profiles of 6 canonical networks. The networks are the default mode network (DMN), the executive control network (ECN), the dorsal attention network (DAN), the salience network (SN), the visual network (VN) and the somatomotor network (SMN) at various axial slices descending posteriorly. All maps are shown in the radiological convention. For reference, approximate anatomical locations for each axial slice are depicted in Fig. S4.

We observed heterogeneous patterns of network connectivity across the subcortex, as cortical networks had distinct mixtures of correlations and anticorrelations within subcortical regions. For example, the ECN showed correlations with the basal ganglia but anticorrelations within the thalamus and medial temporal lobe. These heterogeneities were also seen within individual regions of interest, with many nuclei having diverse patterns of connectivity. For example, the rostral putamen was highly correlated in the ECN and DAN networks, whereas the caudal putamen was anticorrelated. Additionally, we noted that the distribution of connectivity was not consistent across networks, as some networks tended to show more subcortical anticorrelations and some more subcortical correlations. For example, the somatomotor network showed mostly anticorrelations within the subcortex, with only a few small regions of high positive correlations, such as in the amygdala and the ventral posterolateral thalamic nucleus (VPL).

### Correlated and anticorrelated regions

Fig. 3 shows the most correlated subcortical regions for each network. In the final “Combined” column of these panels, the most correlated (thresholded to preserve the top 5% of values for each network) masks were overlapped to visualize regions of network integration.

**Figure 3:**
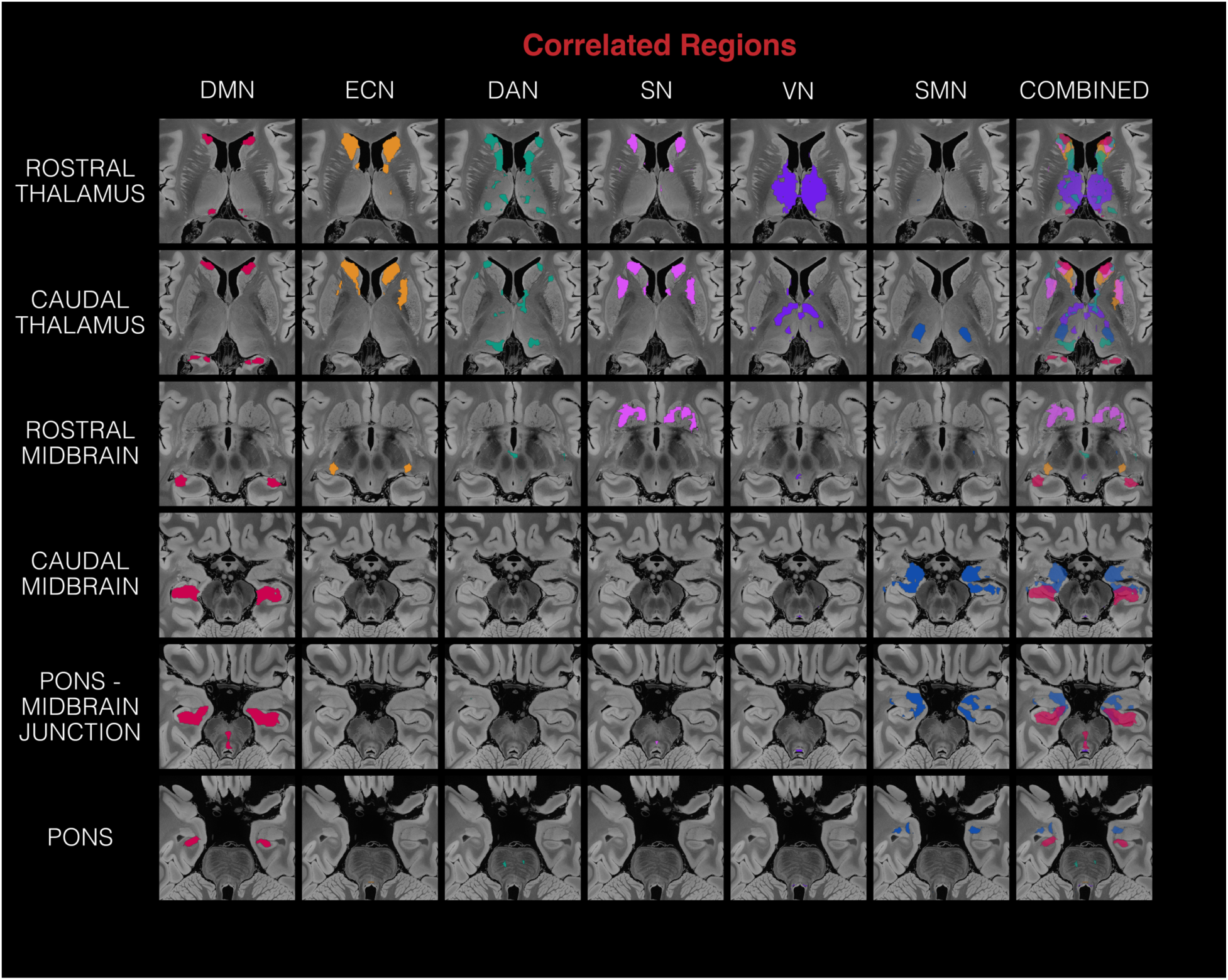
Correlated regions for each network. Each column (except for the last one) represents an individual network, and the colored voxels represent voxels within the top 5% (or the most correlated) of values for that network within the entire subcortex. The last column shows the combination (semi-transparent overlap) of individual network maps.

Fig. 3 shows the regions with highly correlated network overlap within the caudate head and the bed nucleus of the stria terminalis (BNST). Networks that overlapped within the caudate head were the DMN, SN, ECN, and SN. The VN additionally overlapped with the SN, ECN, and the DAN within the BNST and the DAN within the thalamus.

Fig. 4 is the counterpart to Fig. 3, with the thresholding performed to preserve the bottom 5% of values for each network. Fig.4 shows the regions with highly anticorrelated network overlap within the amygdala and the hippocampus. We found that the networks most overlapped within the hippocampus were the DAN, SN, and VN. The DMN, ECN, SN, and VN additionally had a small overlap region within the amygdala. While there were notably few regions of overlap within basal ganglia, thalamic, and brainstem structures, there were slight overlaps between the ECN and SN, as well as the ECN and SMN within the rostral and caudal thalamus.

**Figure 4:**
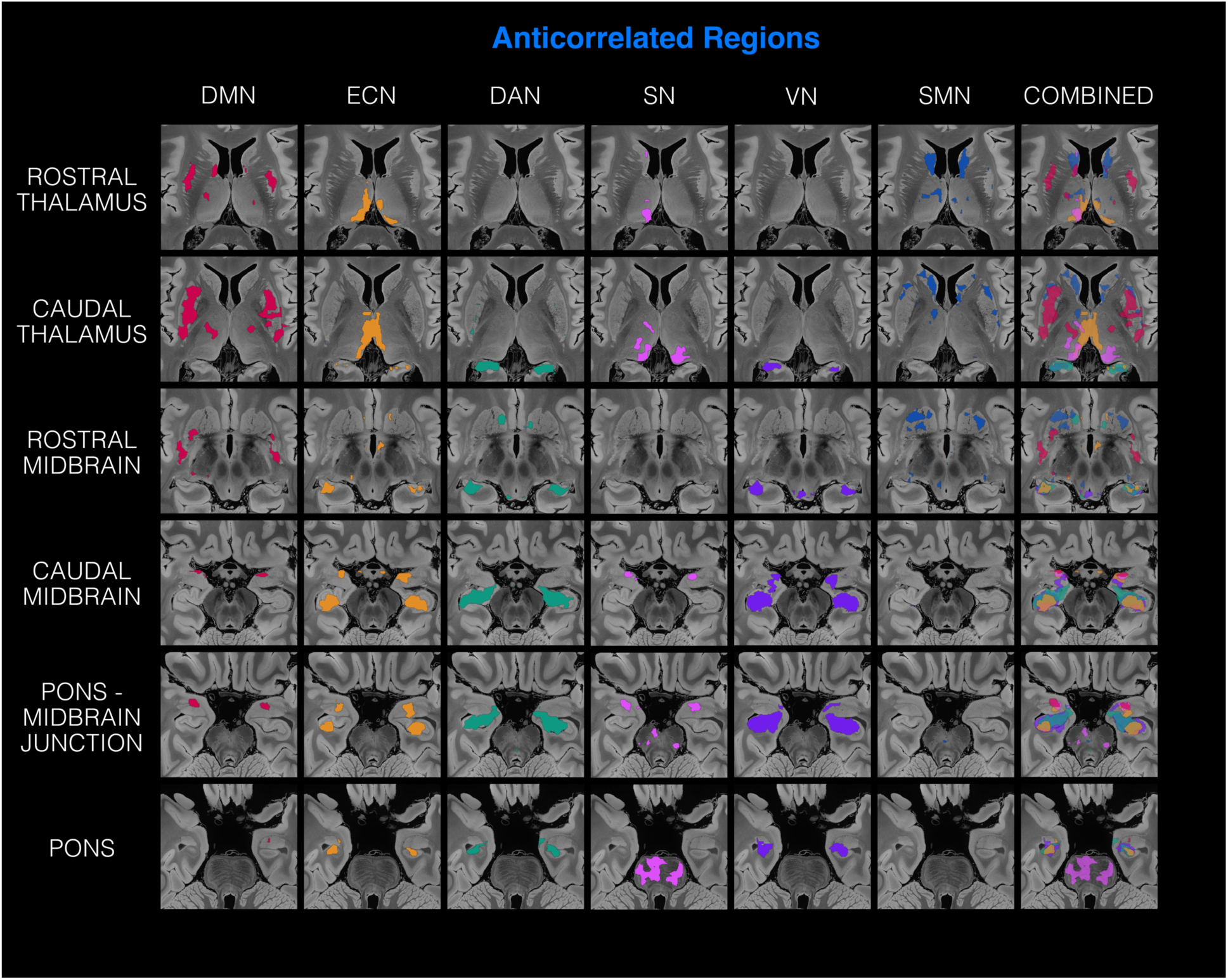
Anticorrelated regions for each network. Each column (except for the last one) represents an individual network, and the colored voxels represent voxels within the bottom 5% (or the most anticorrelated) of values for that network within the entire subcortex. The last column shows the combination (semi-transparent overlap) of individual network maps.

### Correlated, anticorrelated, and mixed network hubs

Regions with multiple network overlaps are shown in Fig. 5, which provides an overview of “correlated network hubs,” “anticorrelated network hubs,” and “mixed network hubs.” In the “correlated” condition (the first pair of columns), the left column defines the number of network overlaps for all six thresholded correlated masks in Fig. 3. For easy reference, the right column shows the combined map from the last column of Fig. 3, indicating the identities of the networks that are overlapping. The middle column pair shows the counterpart for the anticorrelated condition, where the thresholding preserves the bottom 5% of values for each network. For the “mixed network hub” condition in the last column pair, we combined the masks from both the “correlated” and “anticorrelated” conditions.

**Figure 5:**
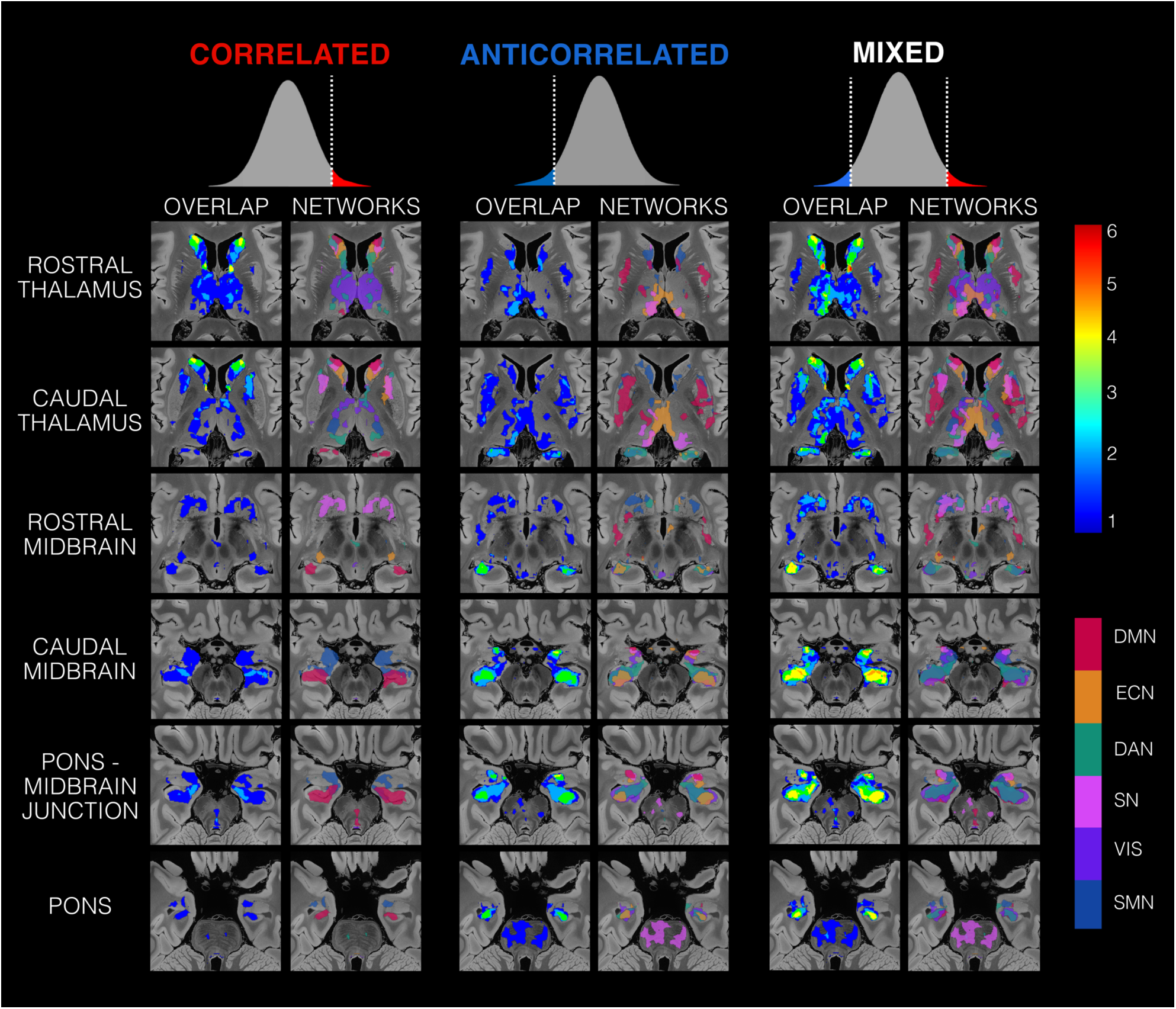
Network overlap and identity maps for the correlated (left column pair), the anticorrelated (middle column pair) and the mixed (right column pair) conditions shown in different axial slices. The left column in each column pair shows the map describing the number of network overlaps at every voxel with values ranging from 0 to 6, as indicated by the top right color bar. The right column in each column pair labels the locations and identity of the thresholded networks overlapping at every voxel, where each color represents a distinct brain network as indicated by the bottom right color bar.

We observed two main regions of correlated network hubs within the caudate head and the bed nucleus of the stria terminalis, where four networks overlapped. The networks overlapping at these two regions were the DMN, ECN, DAN, and SN for the caudate head and the ECN, DAN, SN, and VN for the bed nucleus of the stria terminalis. We also observed two regions of anticorrelated network hubs within the amygdala and the hippocampus, with four and three networks overlapping, respectively. The networks overlapping in these two regions were the DMN, ECN, and SN and VN for the amygdala and the ECN, DAN, and VN for the hippocampus.

When we combined the correlated and anticorrelated network masks to identify mixed network hubs, we identified regions with more heterogeneous relationships to resting-state networks. We identified that a region within the caudate head not only had strong correlated relationships to four higher-order cognitive networks but was also anticorrelated to the somatomotor network. The BNST was notably the only nucleus with strong relationships to all six networks within this “mixed” condition, with a strong correlated relationship to four networks (the ECN, SN, DAN, and VN) and strong anticorrelated relationship to two networks (the DMN, and SMN). A small unilateral region within the amygdala overlap increased to five networks within the mixed condition with the addition of a correlated overlap with the SMN in addition to its previous four anticorrelated network contributions. Within the hippocampus, there was a bilateral region of five network overlaps with the addition of two correlated networks (DMN and SMN) overlapping with three anticorrelated networks (ECN, DAN, VN).

### Regions associated with consciousness level correlate with multiple cortical resting state networks

Figures 6, 7, and 8 describe network hubs within multiple subcortical regions that are associated with consciousness level. Network contour maps were generated for these three figures to identify the highest threshold necessary for there to be 𝑘 networks overlapped at each voxel. 𝑘 = 4 was used to best illustrate the correspondence between the hubs identified in the present work and the hotspots identified in prior studies where lesions are associated with loss of consciousness (Fischer et al., 2016; Parvizi & Damasio, 2003; Snider et al., 2020).

**Figure 6:**
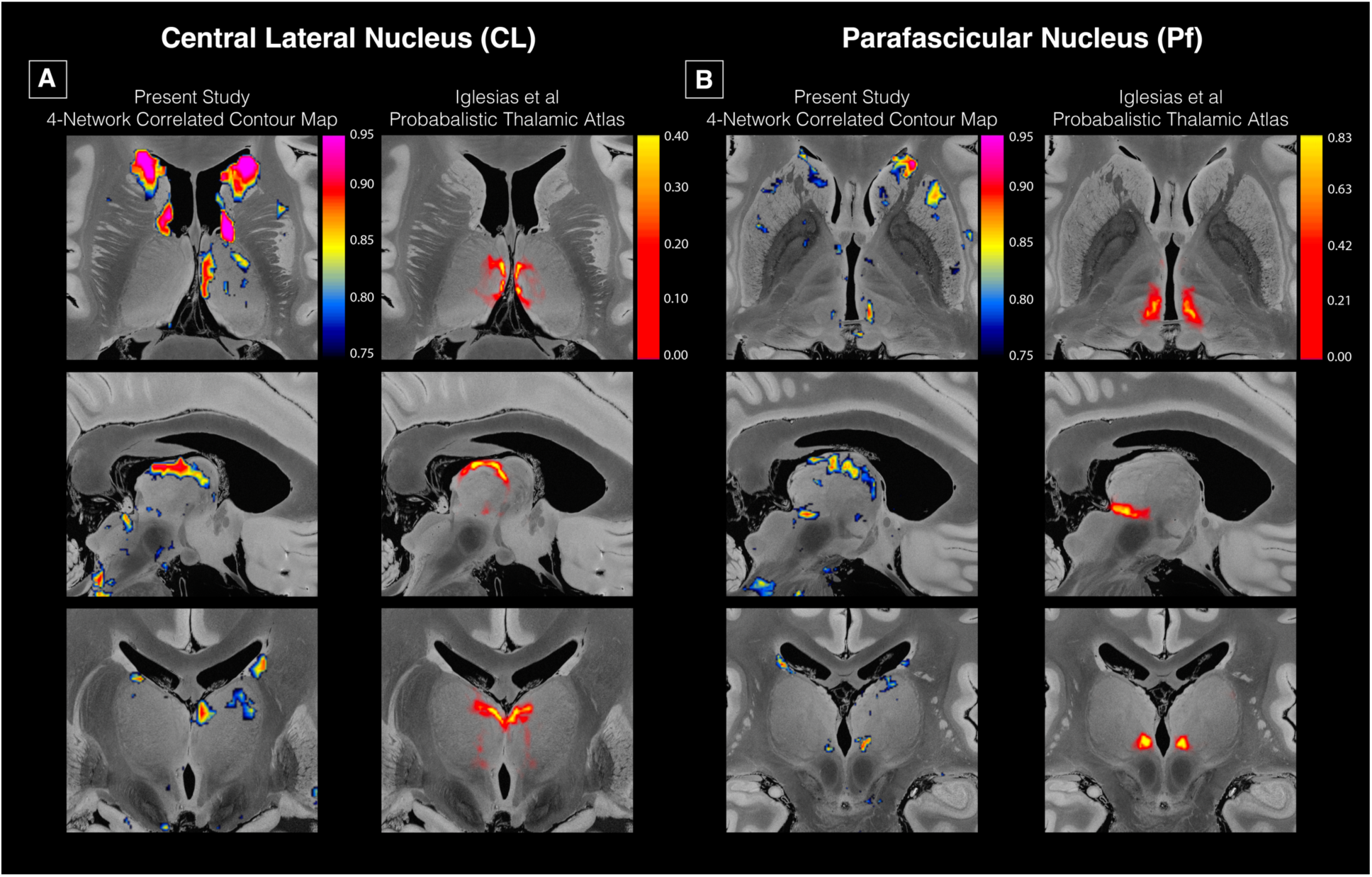
Subcortical correlated connectivity hubs within the central lateral (CL) and parafascicular nuclei (Pf) of the thalamus. (A) A side-by-side comparison of the probabilistic thalamic segmentation atlas (Iglesias et al., 2018) for the central lateral thalamic nucleus alongside the 4-network correlated contour map is shown from an axial (top row) left hemispheric sagittal (middle row) and coronal (bottom row) perspective. (B) The counterpart to (A) but for the Pf.

**Figure 7:**
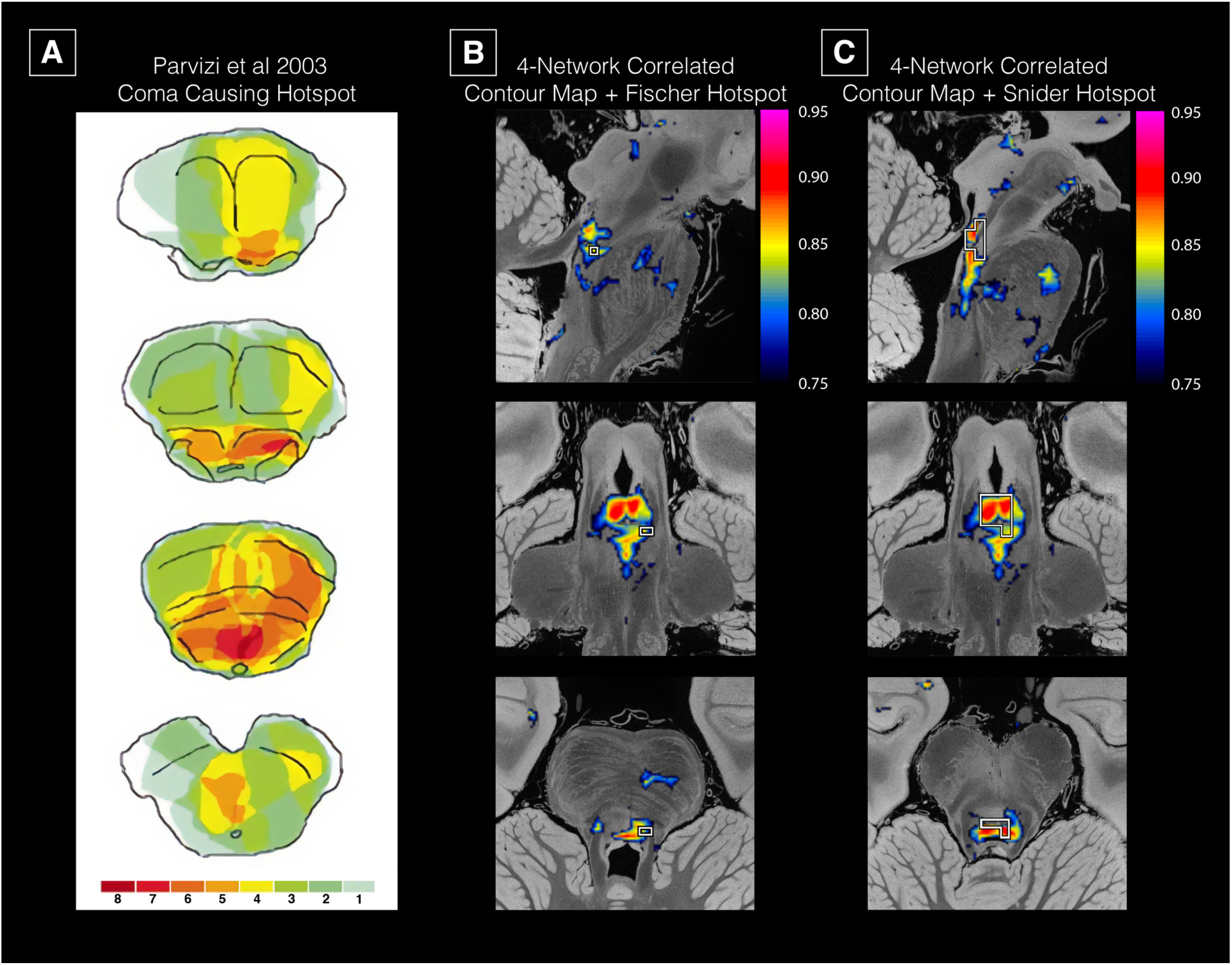
Brainstem tegmentum connectivity hub overlaps with region previously implicated in modulating human consciousness. (A) shows the locations within the brainstem of lesions in patients presenting with coma (figure adapted from Parvizi et al 2003). The colors correspond to the number of patients with lesions in that area. (B) shows our 4-network correlated contour map with a white outlined region labeling the regions of maximum lesion overlap from patients presenting with coma from Fischer et al. 2016. (C) shows our 4-network correlated contour map with a white outlined region labeling another brainstem hotspot whose connectivity to cortical lesions is significantly (p < 0.05) associated with loss of consciousness from Snider et al. 2020.

**Figure 8:**
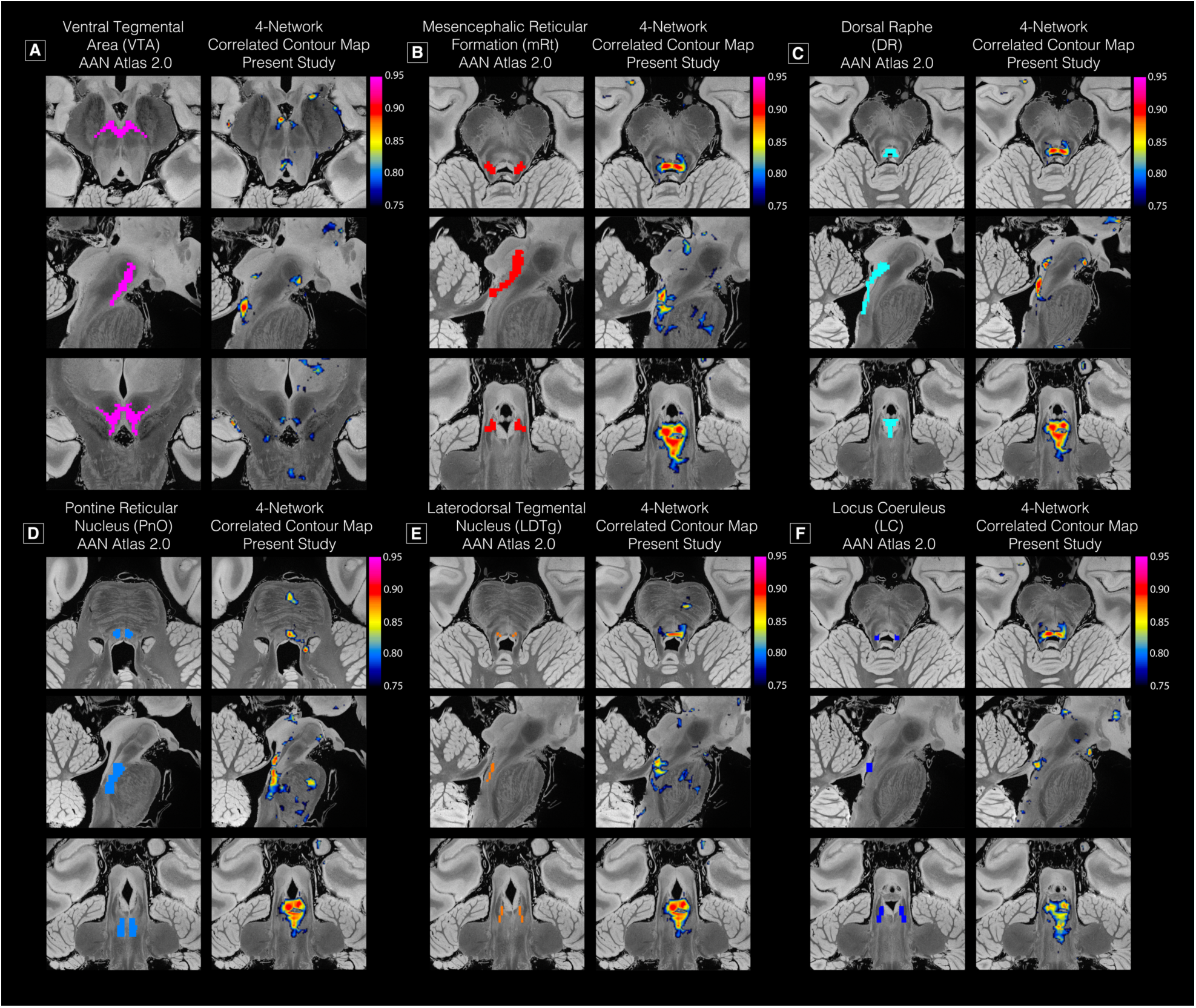
Comparison of our 4-network contour map with the nuclei defined in the Ascending Arousal Network (AAN) atlas. (A) – (E) corresponds to the VTA, mRT, DR, PnO, LDTg, and LC, respectively. Each panel has two columns with axial, sagittal and coronal views of the pons-midbrain junction and pons.

Fig. 6 displays the 4-network correlated contour map findings within the central lateral thalamic nucleus (CL) in (A) left column and for the perifascicular thalamic nucleus (Pf) in (B) left column with the axial, sagittal and coronal views from top to bottom. The right column in both A and B shows the probabilistic values for the CL and Pf nuclei as defined by the probabilistic thalamic segmentation atlas (Iglesias et al., 2018). This distribution of probability values indicates where these nuclei are most likely to be defined in space. The general shape of both the CL and the Pf outlined in this atlas are seen in the 4-network contour map, providing evidence that the CL and the Pf (both intralaminar nuclei) are nuclei within the thalamus with high levels of network integration. For the left hemispheric CL, the networks overlapping were the DMN, DAN, SN, and VN. The Pf finding was also asymmetrical, with a smaller profile on the right hemisphere than on the left. Within the left Pf, the networks overlapping were the DMN, ECN, DAN, and SN. The networks overlapping within the smaller spot in the right Pf were the DMN, DAN, SN, and VS.

Fig. 7 displays network hubs within the brainstem that overlap with three previously reported brainstem “hot spots”: two previously reported regions that cause coma when lesioned (Fischer et al., 2016; Parvizi & Damasio, 2003) and one region whose connectivity to cortical lesions was associated with loss of consciousness (Snider et al., 2020). The first column shows the Parvizi hotspot superimposed on axial sections of the brainstem. The second and third columns show the Fischer and Snider hotspots (white outlines) superimposed on hub connectivity data from the present study. All three previously published hotspots overlap with our 4-network correlated hub within the pontomesencephalic tegmentum. Fig. 8, shows the axial sagittal and coronal slices from the midbrain to the pontine structures of the 4-network contour map alongside six ascending arousal network nuclei as defined by the Harvard Ascending Arousal Network (AAN) Atlas (Edlow et al., 2012). Each panel describes the hotspots in the 4-network contour map, where they overlap with the corresponding AAN nuclei in the left panel. We found that multiple nuclei associated with consciousness level overlap with at least four correlated networks. However, it is notable that there is a consistent pattern of multiple higher-order cognitive networks being positively correlated to multiple structures deemed necessary for sustaining consciousness. Furthermore, this pattern is even more consistent with the salience network, the default mode network, and the visual network, being three of the four correlated networks for all these brainstem nuclei. Anticorrelations and mixed correlations have been observed in other networks. A detailed summary of the relationships between the large-scale resting-state networks and the AAN nuclei is shown in Table 2.

**Table 2:**
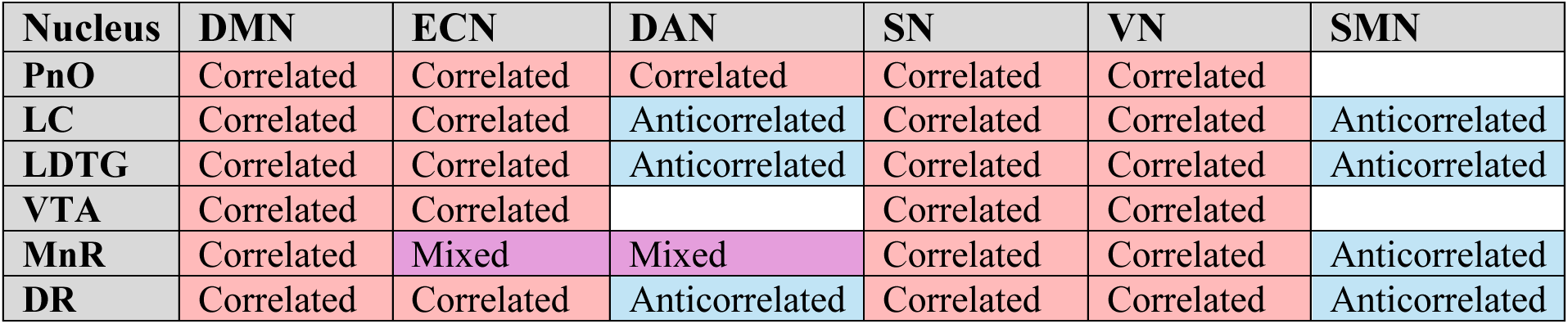
The relationships between large-scale resting state brain networks and AAN nuclei. The “Correlated” relationship indicates that the network’s connectivity values lie in the top 20% of the overall connectivity distribution within the nucleus. The “Anticorrelated” relationship indicates the counterpart for connectivity within the bottom 20% of the distribution. The “Mixed” relationship indicates both correlated and anticorrelated conditions occur within the same nucleus but in different spatial locations. An empty cell indicates no significant correlation was observed.

| Nucleus | DMN | ECN | DAN | SN | VN | SMN |
| --- | --- | --- | --- | --- | --- | --- |
| PnO | Correlated | Correlated | Correlated | Correlated | Correlated |  |
| LC | Correlated | Correlated | Anticorrelated | Correlated | Correlated | Anticorrelated |
| LDTG | Correlated | Correlated | Anticorrelated | Correlated | Correlated | Anticorrelated |
| VTA | Correlated | Correlated |  | Correlated | Correlated |  |
| MnR | Correlated | Mixed | Mixed | Correlated | Correlated | Anticorrelated |
| DR | Correlated | Correlated | Anticorrelated | Correlated | Correlated | Anticorrelated |

## Discussion

This brain mapping study leveraged a 7T rs-fMRI healthy control dataset with 168 subjects and a tensor-based NASCAR decomposition method to map the subcortical connectivity of six canonical resting state networks. We identified multiple subcortical network hubs demonstrating differential patterns of functional connectivity with cortical networks. These hubs included regions that have been historically targeted in therapeutic stimulation studies of patients with DoC: the CL, Pf, and VTA (Chudy et al., 2018; Chudy et al., 2023; Giacino et al., 2012; Schiff et al., 2007). In addition, we found a subcortical connectivity hub in a region of the pontomesencephalic tegmentum that overlaps with multiple reticular and extrareticular arousal nuclei and that has been shown in three prior lesion studies to be a “hot spot” associated with loss of consciousness (Fischer et al., 2016; Parvizi & Damasio, 2003; Snider et al., 2020), strengthening the evidence that this region of the brainstem tegmentum is critical to human consciousness. All brainstem and thalamic hubs were functionally connected to both the DMN and SN, emphasizing the importance of these cortical networks in integrative subcortico-cortical signaling in the human brain. Furthermore, we identified subcortical hubs in the caudate head, putamen, hippocampus, amygdala, and bed nucleus of the stria terminalis – regions that are classically associated with modulation of cognition, behavior, and sensorimotor function. Collectively, these observations provide insights into the subcortical regions of the human brain that are strongly coupled to large-scale cortical networks and identify potential targets for therapeutic neurostimulation studies aimed at promoting recovery of consciousness in patients with DoC.

Our subcortical connectivity findings add to the growing evidence base for a therapeutic paradigm in which subcortical regions are targeted to restore consciousness in patients with DoC. We identified multiple subcortical nodes that were connected to four cortical networks and a small number of subcortical nodes connected to as many as six cortical networks. For example, the caudate head was observed to be a 6-network correlated hub, and the bed nucleus of the stria terminalis a 6-network mixed hub. Accordingly, stimulation of subcortical hub nodes has the potential to activate a broad expanse of cortical neurons across multiple functionally connected networks. In prior clinical trials, electrophysiologic stimulation of the central thalamus (Schiff et al., 2007) and pharmacological stimulation of the VTA (Giacino et al., 2012) have generally yielded more robust behavioral responses in patients with DoC than did transcranial magnetic stimulation or transcranial direct current stimulation (Fan et al., 2022; Thibaut et al., 2014; Wan et al., 2024) of cortical nodes. Our connectivity maps provide a potential mechanistic basis for these clinical trial results, as the CL and VTA were each found to be correlated to 4 cortical networks: DMN, SN, DAN, and VN in the CL, and DMN, FPN, SN, and VN in the VTA.

Notably, while a subcortical region of 4-network correlated activity showed substantial spatial overlap with the probabilistic definition of the CL using the Iglesias atlas (Fig. 5), this finding was only found in the left CL. We propose that this asymmetry is due to hemispheric heterogeneities in the SN, whereby the left thalamus has significantly higher connectivity values than in the right thalamus (P < 0.001 Mann Whitney U Test). Interestingly, the correlated hub in the left CL additionally had a subregion of 2-network anticorrelated overlap, meaning that part of the CL represented a 6-network mixed hub (Fig. S2). While the VTA’s 4-network correlated hub did not overlap with additional anticorrelated networks, there were regions within the VTA that overlapped with a 5- network mixed hub consisting of three correlated networks and two anticorrelated networks (Fig. S3). These observations reveal highly specialized subregions of subcortical hubs that may differentially modulate cortical networks via activating and inhibitory signaling.

The physiologic meaning of rs-fMRI anticorrelations within functional networks continues to be debated (Fox et al., 2005; Kevin & Michael, 2017), and the microscale synaptic correlates of mesoscale anticorrelations remain unclear (DeFelipe, 2010). Elucidating the therapeutic potential of a correlated hub versus that of a mixed hub, therefore, requires further inquiry in studies that link mesoscale neuroimaging with microscale electrophysiology measures. Nevertheless, our functional connectivity maps are consistent with electrophysiologic studies showing a “funnel effect” of information processing in the human brain, whereby information dimensionality is reduced within subcortical structures (Blouw et al., 2016; Bota et al., 2015) such that stimulating a small hub node in the subcortex generates widespread activation of the cortex (Horn et al., 2017; Horn et al., 2019). By beginning to link rs-fMRI connectivity maps to prior patterns observed in electrophysiologic data, our findings highlight the potential for rs-fMRI to serve as a clinically relevant tool that can identify subcortical therapeutic targets in patients with DoC.

The functional connectivity maps generated here, which we release to the academic community, expand the repertoire of therapeutic targets that can be considered in future clinical trials. While the central thalamus has been the most common target of electrophysiologic and ultrasonic therapies, and while the VTA has been the most common target of pharmacologic therapies, our findings suggest that hub nodes in the basal ganglia, medial temporal lobe, and bed nucleus of the stria terminalis warrant future consideration as therapeutic targets in studies that aim to modulate cognition, behavior, and sensorimotor function in patients with severe brain injuries. Ultrasonic stimulation studies in patients with DoC have begun to target the basal ganglia (Cain, Visagan, et al., 2021) based on the mesocircuit hypothesis of consciousness (Schiff, 2010), in which a striato-pallido-thalamic circuit is postulated to modulate a broad expanse of fronto-parietal cortex. The mechanistic relationships of the caudate and putamen hubs identified here to the mesocircuit is beyond the scope of the present work, but our findings add to growing evidence that it may be possible to identify the functional integrity of the mesocircuit using ultra-high resolution rs-fMRI (Li, Curley, et al., 2021), an advance that would have substantial implications for personalized therapy selection in patients with DoC who have a disrupted mesocircuit.

Although mesoscale measurements of functional connectivity are only surrogate markers of microscale signaling at the synaptic level, our identification of correlated, anticorrelated, and mixed hubs sheds new light on how subcortical ensembles of neurons may differentially modulate cortical networks. We found that subcortical hubs in the brainstem tegmentum and central thalamus had high levels of correlated connectivity with four or more cortical networks. These regions were consistently correlated with the DMN, SN, and VN, anticorrelated with the SMN, and had mixed connectivity properties with the DAN and ECN. In contrast, the bed nucleus of the stria terminalis contained high levels of network overlap but with varying degrees (top 5% for SN, FPN, VN, DAN, and bottom 5% for the DMN and SMN) representing a mixed hub node that has the highest level of correlated activity from four networks and the highest level of anticorrelated activity from two networks. A region within the anterior putamen had the same mixed type of integration pattern as the bed nucleus of the stria terminalis (correlated to the SN, FPN, VN, DAN, and anticorrelated to SMN and DMN) but with a lower threshold. Future studies combining rs-fMRI and intracranial electrophysiologic recordings are needed to elucidate the biological basis of mixed subcortical hubs (Stieger et al., 2024). We postulate that a subcortical node correlated with four task-positive networks and anticorrelated with the task-negative DMN may contribute to the brain’s ability to “toggle” between task-positive and task-negative (i.e., resting) states.

Our observations about functional connectivity hubs in the healthy, conscious human brain also provide a mechanistic basis for previous descriptions of subcortical lesions that cause coma in patients with severe brain injuries. Specifically, we identified a subcortical hub that includes multiple reticular and extrareticular arousal nuclei and that is located in the same “hot spot” region of the pontomesencephalic tegmentum that was found in prior lesion studies to cause coma (Fischer et al., 2016; Parvizi & Damasio, 2003) or to be associated with loss of consciousness (Snider et al., 2020). While the pathophysiologic mechanism of coma onset in patients with brainstem lesions continues to be debated, our functional connectivity results indicate that injury to a subcortical hub causes deafferentation and downregulation of cortical networks, such as the DMN, that are essential for consciousness. Given that recovery of consciousness after severe brain injury is associated with the reemergence of multiple cortical networks (Demertzi et al., 2015), a key area for future inquiry will be to identify the combinations of subcortical hubs and their associated cortical networks that are necessary and sufficient for recovery of consciousness.

Several limitations should be considered when interpreting these results in the context of future clinical trials that aim to restore consciousness after severe brain injury. While some subcortical hubs identified here have been previously associated with consciousness, we also identified widely connected subcortical hubs in the basal ganglia, medial temporal lobe, and bed nucleus of the stria terminalis that are historically associated with cognition, behavior, and sensorimotor function, not conscious awareness. For example, we found high levels of network overlap bilaterally within the caudate head for four networks when thresholding to include the top 2% of values, five networks when thresholding to the top 10%, and even a bilateral cluster of voxels of six network correlated overlap when thresholding to the top 15% of values. While lesions within the caudate can cause cognitive and behavioral dysfunction (Bokura & Robinson, 1997; Graff-Radford et al., 2017; Mendez et al., 1989), to the best of our knowledge, there have been no associations between caudate lesions and loss of consciousness. For this reason, it is important to consider that the identification of a widely connected subcortical hub via rs-fMRI data does not prove that the hub modulates consciousness. Rather, these connectivity data must be interpreted in the context of prior animal and human studies that have linked each subcortical hub to distinct aspects of consciousness, cognition, behavior, or sensorimotor function.

A methodological limitation of the present work is that identification of subcortical hubs at the group level requires precise registration across subjects, which was enabled here by the NASCAR network identification pipeline (Li, Curley, et al., 2021). Recent work based on deep neural network methods have been shown to improve inter-subject registration (Balakrishnan et al., 2019; Cheng et al., 2020), but registration of subcortical structures remains challenging due to the low resolution and low SNR of the fMRI data, as well as the potential mismatch between human brain anatomy and function (Li et al., 2024). Therefore, inferences should not be made on any results reported here at voxel resolution. For instance, Fig. 3 illustrates hubs that are consistently located within the hippocampus across multiple networks. The small overlap in the amygdala, although also across multiple networks, may not provide sufficiently strong evidence of hubness because of potentially inaccurate inter-subject registration.

In conclusion, we identified subcortical connectivity hubs in the VTA, CL, and Pf – regions that have historically been targeted in therapeutic trials to restore consciousness in patients with severe brain injuries. An additional hub within the pontomesencephalic tegmentum overlapped with a previously described coma-causing “hot spot,” indicating that this brainstem region is a potential therapeutic target for neuromodulation of consciousness. Subcortical hubs were also identified in regions of the caudate, putamen, medial temporal lobe, and bed nucleus of the stria terminalis that are believed to modulate cognition, behavior and sensorimotor function. While further evidence from multimodal neuroimaging and electrophysiologic studies is needed to elucidate the specific functional role of each subcortical hub, these results strengthen the evidence base for targeting subcortical hubs as a therapeutic strategy to restore consciousness in patients with DoC.

## Author Contributions

BLE, JL, and MKC conceived and designed the study. MKC, JL, and BLE wrote the original draft of the manuscript. MKC and JL processed and analyzed the data. AH and LL contributed to the study design and interpreted the data. BLE and JL contributed to project administration and supervision. All authors contributed to editing the manuscript.

## Data Availability

The data used in this study are publicly available from the Wash U/U Minn component of the Human Connectome Project, Young Adult Study at https://www.humanconnectome.org/study/hcp-young-adult. For research purposes, we release the code at the GitHub repository (https://github.com/ComaRecoveryLab/Subcortical_Network_Hubs).

## Acknowledgements

The authors thank Drs. David Fischer and Samuel B. Snider for providing the neuroanatomic coordinates of their brainstem hotspots for comparison to the results in the present study.

## References

1. Akrami, H., Joshi, A., Li, J., & Leahy, R. (2019). Group-wise alignment of resting fMRI in space and time (Vol. 10949). SPIE. 10.1117/12.2512564

2. Anand, A. J., Minqi, C., Jian, L., Soyoung, C., & Richard, M. L. (2018). Are you thinking what I’m thinking? Synchronization of resting fMRI time-series across subjects. Neuroimage, 172, 740–752. 10.1016/j.neuroimage.2018.01.058

3. Aston-Jones, G., Chen, S., Zhu, Y., & Oshinsky, M. L. (2001). A neural circuit for circadian regulation of arousal. Nat Neurosci, 4(7), 732–738. 10.1038/89522

4. Balakrishnan, G., Zhao, A., Sabuncu, M. R., Guttag, J., & Dalca, A. V. (2019). VoxelMorph: A Learning Framework for Deformable Medical Image Registration. IEEE Trans Med Imaging. 10.1109/TMI.2019.2897538

5. Blouw, P., Solodkin, E., Thagard, P., & Eliasmith, C. (2016). Concepts as Semantic Pointers: A Framework and Computational Model. Cognitive Science, 40(5), 1128–1162. 10.1111/cogs.12265

6. Bokura, H., & Robinson, R. G. (1997). Long-term cognitive impairment associated with caudate stroke. Stroke, 28(5), 970–975. 10.1161/01.str.28.5.970

7. Bota, M., Sporns, O., & Swanson, L. W. (2015). Architecture of the cerebral cortical association connectome underlying cognition. Proc Natl Acad Sci U S A, 112(16), E2093–2101. 10.1073/pnas.1504394112

8. Cain, J. A., Spivak, N. M., Coetzee, J. P., Crone, J. S., Johnson, M. A., Lutkenhoff, E. S., Real, C., Buitrago-Blanco, M., Vespa, P. M., Schnakers, C., & Monti, M. M. (2021). Ultrasonic thalamic stimulation in chronic disorders of consciousness. Brain Stimul, 14(2), 301–303. 10.1016/j.brs.2021.01.008

9. Cain, J. A., Spivak, N. M., Coetzee, J. P., Crone, J. S., Johnson, M. A., Lutkenhoff, E. S., Real, C., Buitrago-Blanco, M., Vespa, P. M., Schnakers, C., & Monti, M. M. (2022). Ultrasonic Deep Brain Neuromodulation in Acute Disorders of Consciousness: A Proof-of-Concept. Brain Sci, 12(4). 10.3390/brainsci12040428

10. Cain, J. A., Visagan, S., Johnson, M. A., Crone, J., Blades, R., Spivak, N. M., Shattuck, D. W., & Monti, M. M. (2021). Real time and delayed effects of subcortical low intensity focused ultrasound. Sci Rep, 11(1), 6100. 10.1038/s41598-021-85504-y

11. Cheng, J., Dalca, A. V., Fischl, B., Zollei, L., & Alzheimer’s Disease Neuroimaging, I. (2020). Cortical surface registration using unsupervised learning. Neuroimage, 221, 117161. 10.1016/j.neuroimage.2020.117161

12. Chudy, D., Deletis, V., Almahariq, F., Marcinkovic, P., Skrlin, J., & Paradzik, V. (2018). Deep brain stimulation for the early treatment of the minimally conscious state and vegetative state: experience in 14 patients. J Neurosurg, 128(4), 1189–1198. 10.3171/2016.10.JNS161071

13. Chudy, D., Deletis, V., Paradzik, V., Dubroja, I., Marcinkovic, P., Oreskovic, D., Chudy, H., & Raguz, M. (2023). Deep brain stimulation in disorders of consciousness: 10 years of a single center experience. Sci Rep, 13(1), 19491. 10.1038/s41598-023-46300-y

14. Crossley, N. A., Mechelli, A., Scott, J., Carletti, F., Fox, P. T., McGuire, P., & Bullmore, E. T. (2014). The hubs of the human connectome are generally implicated in the anatomy of brain disorders. Brain, 137(Pt 8), 2382–2395. 10.1093/brain/awu132

15. DeFelipe, J. (2010). From the connectome to the synaptome: an epic love story. Science, 330(6008), 1198–1201. 10.1126/science.1193378

16. Demertzi, A., Antonopoulos, G., Heine, L., Voss, H. U., Crone, J. S., de Los Angeles, C., Bahri, M. A., Di Perri, C., Vanhaudenhuyse, A., Charland-Verville, V., Kronbichler, M., Trinka, E., Phillips, C., Gomez, F., Tshibanda, L., Soddu, A., Schiff, N. D., Whitfield-Gabrieli, S., & Laureys, S. (2015). Intrinsic functional connectivity differentiates minimally conscious from unresponsive patients. Brain, 138(Pt 9), 2619–2631. 10.1093/brain/awv169

17. Edlow, B. L., Barra, M. E., Zhou, D. W., Foulkes, A. S., Snider, S. B., Threlkeld, Z. D., Chakravarty, S., Kirsch, J. E., Chan, S. T., Meisler, S. L., Bleck, T. P., Fins, J. J., Giacino, J. T., Hochberg, L. R., Solt, K., Brown, E. N., & Bodien, Y. G. (2020). Personalized Connectome Mapping to Guide Targeted Therapy and Promote Recovery of Consciousness in the Intensive Care Unit. Neurocrit Care, 33(2), 364–375. 10.1007/s12028-020-01062-7

18. Edlow, B. L., Claassen, J., Schiff, N. D., & Greer, D. M. (2021). Recovery from disorders of consciousness: mechanisms, prognosis and emerging therapies. Nat Rev Neurol, 17(3), 135–156. 10.1038/s41582-020-00428-x

19. Edlow, B. L., Haynes, R. L., Takahashi, E., Klein, J. P., Cummings, P., Benner, T., Greer, D. M., Greenberg, S. M., Wu, O., Kinney, H. C., & Folkerth, R. D. (2013). Disconnection of the ascending arousal system in traumatic coma. J Neuropathol Exp Neurol, 72(6), 505–523. 10.1097/NEN.0b013e3182945bf6

20. Edlow, B. L., Olchanyi, M., Freeman, H. J., Li, J., Maffei, C., Snider, S. B., Zollei, L., Iglesias, J. E., Augustinack, J., Bodien, Y. G., Haynes, R. L., Greve, D. N., Diamond, B. R., Stevens, A., Giacino, J. T., Destrieux, C., van der Kouwe, A., Brown, E. N., Folkerth, R. D., . . . Kinney, H. C. (2024). Multimodal MRI reveals brainstem connections that sustain wakefulness in human consciousness. Sci Transl Med, 16(745), eadj4303. 10.1126/scitranslmed.adj4303

21. Edlow, B. L., Sanz, L. R. D., Polizzotto, L., Pouratian, N., Rolston, J. D., Snider, S. B., Thibaut, A., Stevens, R. D., Gosseries, O., Curing Coma, C., & its contributing, m. (2021). Therapies to Restore Consciousness in Patients with Severe Brain Injuries: A Gap Analysis and Future Directions. Neurocrit Care, 35(Suppl 1), 68–85. 10.1007/s12028-021-01227-y

22. Edlow, B. L., Takahashi, E., Wu, O., Benner, T., Dai, G., Bu, L., Grant, P. E., Greer, D. M., Greenberg, S. M., Kinney, H. C., & Folkerth, R. D. (2012). Neuroanatomic connectivity of the human ascending arousal system critical to consciousness and its disorders. J Neuropathol Exp Neurol, 71(6), 531–546. 10.1097/NEN.0b013e3182588293

23. Elias, G. J. B., Loh, A., Gwun, D., Pancholi, A., Boutet, A., Neudorfer, C., Germann, J., Namasivayam, A., Gramer, R., Paff, M., & Lozano, A. M. (2021). Deep brain stimulation of the brainstem. Brain, 144(3), 712–723. 10.1093/brain/awaa374

24. Fan, J., Zhong, Y., Wang, H., Aierken, N., & He, R. (2022). Repetitive transcranial magnetic stimulation improves consciousness in some patients with disorders of consciousness. Clin Rehabil, 36(7), 916–925. 10.1177/02692155221089455

25. Fischer, D. B., Boes, A. D., Demertzi, A., Evrard, H. C., Laureys, S., Edlow, B. L., Liu, H., Saper, C. B., Pascual-Leone, A., Fox, M. D., & Geerling, J. C. (2016). A human brain network derived from coma-causing brainstem lesions. Neurology, 87(23), 2427–2434. 10.1212/WNL.0000000000003404

26. Fox, M. D., Snyder, A. Z., Vincent, J. L., Corbetta, M., Van Essen, D. C., & Raichle, M. E. (2005). The human brain is intrinsically organized into dynamic, anticorrelated functional networks. Proc Natl Acad Sci U S A, 102(27), 9673–9678. 10.1073/pnas.0504136102

27. Fridman, E. A., Osborne, J. R., Mozley, P. D., Victor, J. D., & Schiff, N. D. (2019). Presynaptic dopamine deficit in minimally conscious state patients following traumatic brain injury. Brain, 142(7), 1887–1893. 10.1093/brain/awz118

28. Fuller, P. M., Sherman, D., Pedersen, N. P., Saper, C. B., & Lu, J. (2011). Reassessment of the structural basis of the ascending arousal system. J Comp Neurol, 519(5), 933–956. 10.1002/cne.22559

29. Giacino, J. T., Whyte, J., Bagiella, E., Kalmar, K., Childs, N., Khademi, A., Eifert, B., Long, D., Katz, D. I., Cho, S., Yablon, S. A., Luther, M., Hammond, F. M., Nordenbo, A., Novak, P., Mercer, W., Maurer-Karattup, P., & Sherer, M. (2012). Placebo-controlled trial of amantadine for severe traumatic brain injury. N Engl J Med, 366(9), 819–826. 10.1056/NEJMoa1102609

30. Girn, M., Setton, R., Turner, G. R., & Spreng, R. N. (2024). The “limbic network,” comprising orbitofrontal and anterior temporal cortex, is part of an extended default network: Evidence from multi-echo fMRI. Network Neuroscience, 8(3), 860–882. 10.1162/netn_a_00385

31. Glasser, M. F., Smith, S. M., Marcus, D. S., Andersson, J. L., Auerbach, E. J., Behrens, T. E., Coalson, T. S., Harms, M. P., Jenkinson, M., Moeller, S., Robinson, E. C., Sotiropoulos, S. N., Xu, J., Yacoub, E., Ugurbil, K., & Van Essen, D. C. (2016). The Human Connectome Project’s neuroimaging approach. Nat Neurosci, 19(9), 1175–1187. 10.1038/nn.4361

32. Glasser, M. F., Sotiropoulos, S. N., Wilson, J. A., Coalson, T. S., Fischl, B., Andersson, J. L., Xu, J., Jbabdi, S., Webster, M., Polimeni, J. R., Van Essen, D. C., Jenkinson, M., & Consortium, W. U.-M. H. (2013). The minimal preprocessing pipelines for the Human Connectome Project. Neuroimage, 80, 105–124. 10.1016/j.neuroimage.2013.04.127

33. Graff-Radford, J., Williams, L., Jones, D. T., & Benarroch, E. E. (2017). Caudate nucleus as a component of networks controlling behavior. Neurology, 89(21), 2192–2197. doi:10.1212/WNL.0000000000004680

34. Horn, A., & Fox, M. D. (2020). Opportunities of connectomic neuromodulation. Neuroimage, 221, 117180. 10.1016/j.neuroimage.2020.117180

35. Horn, A., Reich, M., Vorwerk, J., Li, N., Wenzel, G., Fang, Q., Schmitz-Hubsch, T., Nickl, R., Kupsch, A., Volkmann, J., Kuhn, A. A., & Fox, M. D. (2017). Connectivity Predicts deep brain stimulation outcome in Parkinson disease. Ann Neurol, 82(1), 67–78. 10.1002/ana.24974

36. Horn, A., Wenzel, G., Irmen, F., Huebl, J., Li, N., Neumann, W. J., Krause, P., Bohner, G., Scheel, M., & Kuhn, A. A. (2019). Deep brain stimulation induced normalization of the human functional connectome in Parkinson’s disease. Brain, 142(10), 3129–3143. 10.1093/brain/awz239

37. Iglesias, J. E., Insausti, R., Lerma-Usabiaga, G., Bocchetta, M., Van Leemput, K., Greve, D. N., van der Kouwe, A., Fischl, B., Caballero-Gaudes, C., & Paz-Alonso, P. M. (2018). A probabilistic atlas of the human thalamic nuclei combining ex vivo MRI and histology. Neuroimage, 183, 314–326. 10.1016/j.neuroimage.2018.08.012

38. Kevin, M., & Michael, D. F. (2017). Towards a consensus regarding global signal regression for resting state functional connectivity MRI. Neuroimage, 154, 169–173. 10.1016/j.neuroimage.2016.11.052

39. Koch, C., Massimini, M., Boly, M., & Tononi, G. (2016). Neural correlates of consciousness: progress and problems. Nat Rev Neurosci, 17(5), 307–321. 10.1038/nrn.2016.22

40. Koike, S., Tanaka, S. C., Okada, T., Aso, T., Yamashita, A., Yamashita, O., Asano, M., Maikusa, N., Morita, K., Okada, N., Fukunaga, M., Uematsu, A., Togo, H., Miyazaki, A., Murata, K., Urushibata, Y., Autio, J., Ose, T., Yoshimoto, J., . . . Brain, M. B. H. B. M. R. I. G. (2021). Brain/MINDS beyond human brain MRI project: A protocol for multi-level harmonization across brain disorders throughout the lifespan. Neuroimage Clin, 30, 102600. 10.1016/j.nicl.2021.102600

41. Lee, M. H., Smyser, C. D., & Shimony, J. S. (2013). Resting-state fMRI: a review of methods and clinical applications. AJNR Am J Neuroradiol, 34(10), 1866–1872. 10.3174/ajnr.A3263

42. Lemon, R. N., & Edgley, S. A. (2010). Life without a cerebellum. Brain, 133(Pt 3), 652–654. 10.1093/brain/awq030

43. Li, J., Choi, S., Joshi, A. A., Wisnowski, J. L., & Leahy, R. M. (2018). Global Pdf-Based Temporal Non-Local Means Filtering Reveals Individual Differences in Brain Connectivity. Proc IEEE Int Symp Biomed Imaging, 2018, 15–19. 10.1109/ISBI.2018.8363513

44. Li, J., Curley, W. H., Guerin, B., Dougherty, D. D., Dalca, A. V., Fischl, B., Horn, A., & Edlow, B. L. (2021). Mapping the subcortical connectivity of the human default mode network. Neuroimage, 245, 118758. 10.1016/j.neuroimage.2021.118758

45. Li, J., Liu, Y., Wisnowski, J. L., & Leahy, R. M. (2023). Identification of overlapping and interacting networks reveals intrinsic spatiotemporal organization of the human brain. Neuroimage, 270, 119944. 10.1016/j.neuroimage.2023.119944

46. Li, J., Tuckute, G., Fedorenko, E., Edlow, B. L., Dalca, A. V., & Fischl, B. (2024). JOSA: Joint surface-based registration and atlas construction of brain geometry and function. Med Image Anal, 98, 103292. 10.1016/j.media.2024.103292

47. Li, J., Wisnowski, J. L., Joshi, A. A., & Leahy, R. M. (2021). Robust brain network identification from multi-subject asynchronous fMRI data. Neuroimage, 227, 117615. 10.1016/j.neuroimage.2020.117615

48. Luppi, A. I., Gellersen, H. M., Liu, Z. Q., Peattie, A. R. D., Manktelow, A. E., Adapa, R., Owen, A. M., Naci, L., Menon, D. K., Dimitriadis, S. I., & Stamatakis, E. A. (2024). Systematic evaluation of fMRI data-processing pipelines for consistent functional connectomics. Nat Commun, 15(1), 4745. 10.1038/s41467-024-48781-5

49. Mendez, M. F., Adams, N. L., & Lewandowski, K. S. (1989). Neurobehavioral changes associated with caudate lesions. Neurology, 39(3), 349–354. 10.1212/wnl.39.3.349

50. Moruzzi, G., & Magoun, H. W. (1949). Brain stem reticular formation and activation of the EEG. Electroencephalogr Clin Neurophysiol, 1(4), 455–473. https://www.ncbi.nlm.nih.gov/pubmed/18421835

51. Pais-Roldan, P., Edlow, B. L., Jiang, Y., Stelzer, J., Zou, M., & Yu, X. (2019). Multimodal assessment of recovery from coma in a rat model of diffuse brainstem tegmentum injury. Neuroimage, 189, 615–630. 10.1016/j.neuroimage.2019.01.060

52. Parvizi, J., & Damasio, A. R. (2003). Neuroanatomical correlates of brainstem coma. Brain, 126(Pt 7), 1524–1536. 10.1093/brain/awg166

53. Raichle, M. E. (2011). The restless brain. Brain Connect, 1(1), 3–12. 10.1089/brain.2011.0019

54. Redinbaugh, M. J., Phillips, J. M., Kambi, N. A., Mohanta, S., Andryk, S., Dooley, G. L., Afrasiabi, M., Raz, A., & Saalmann, Y. B. (2020). Thalamus Modulates Consciousness via Layer-Specific Control of Cortex. Neuron, 106(1), 66–75 e12. 10.1016/j.neuron.2020.01.005

55. Scammell, T. E., Arrigoni, E., & Lipton, J. O. (2017). Neural Circuitry of Wakefulness and Sleep. Neuron, 93(4), 747–765. 10.1016/j.neuron.2017.01.014

56. Schiff, N. D. (2010). Recovery of consciousness after brain injury: a mesocircuit hypothesis. Trends Neurosci, 33(1), 1–9. 10.1016/j.tins.2009.11.002

57. Schiff, N. D., Giacino, J. T., Kalmar, K., Victor, J. D., Baker, K., Gerber, M., Fritz, B., Eisenberg, B., Biondi, T., O’Connor, J., Kobylarz, E. J., Farris, S., Machado, A., McCagg, C., Plum, F., Fins, J. J., & Rezai, A. R. (2007). Behavioural improvements with thalamic stimulation after severe traumatic brain injury. Nature, 448(7153), 600–603. 10.1038/nature06041

58. Smith, S. M., Beckmann, C. F., Andersson, J., Auerbach, E. J., Bijsterbosch, J., Douaud, G., Duff, E., Feinberg, D. A., Griffanti, L., Harms, M. P., Kelly, M., Laumann, T., Miller, K. L., Moeller, S., Petersen, S., Power, J., Salimi-Khorshidi, G., Snyder, A. Z., Vu, A. T., . . . Consortium, W. U.-M. H. (2013). Resting-state fMRI in the Human Connectome Project. Neuroimage, 80, 144–168. 10.1016/j.neuroimage.2013.05.039

59. Snider, S. B., Hsu, J., Darby, R. R., Cooke, D., Fischer, D., Cohen, A. L., Grafman, J. H., & Fox, M. D. (2020). Cortical lesions causing loss of consciousness are anticorrelated with the dorsal brainstem. Hum Brain Mapp, 41(6), 1520–1531. 10.1002/hbm.24892

60. Steriade, M., & Glenn, L. L. (1982). Neocortical and caudate projections of intralaminar thalamic neurons and their synaptic excitation from midbrain reticular core. J Neurophysiol, 48(2), 352–371. 10.1152/jn.1982.48.2.352

61. Stieger, J. R., Pinheiro-Chagas, P., Fang, Y., Li, J., Lusk, Z., Perry, C. M., Girn, M., Contreras, D., Chen, Q., Huguenard, J. R., Spreng, R. N., Edlow, B. L., Wagner, A. D., Buch, V., & Parvizi, J. (2024). Cross-regional coordination of activity in the human brain during autobiographical self-referential processing. Proc Natl Acad Sci U S A, 121(32), e2316021121. 10.1073/pnas.2316021121

62. Tasserie, J., Uhrig, L., Sitt, J. D., Manasova, D., Dupont, M., Dehaene, S., & Jarraya, B. (2022). Deep brain stimulation of the thalamus restores signatures of consciousness in a nonhuman primate model. Sci Adv, 8(11), eabl5547. 10.1126/sciadv.abl5547

63. Thibaut, A., Bruno, M. A., Ledoux, D., Demertzi, A., & Laureys, S. (2014). tDCS in patients with disorders of consciousness: sham-controlled randomized double-blind study. Neurology, 82(13), 1112–1118. 10.1212/WNL.0000000000000260

64. van den Heuvel, M. P., & Sporns, O. (2013). Network hubs in the human brain. Trends Cogn Sci, 17(12), 683–696. 10.1016/j.tics.2013.09.012

65. Vertes, R. P., & Martin, G. F. (1988). Autoradiographic analysis of ascending projections from the pontine and mesencephalic reticular formation and the median raphe nucleus in the rat. J Comp Neurol, 275(4), 511–541. 10.1002/cne.902750404

66. Wan, X., Zhang, Y., Li, Y., & Song, W. (2024). Effects of parietal repetitive transcranial magnetic stimulation in prolonged disorders of consciousness: A pilot study. Heliyon, 10(9), e30192. 10.1016/j.heliyon.2024.e30192

67. Whyte, J., Rajan, R., Rosenbaum, A., Katz, D., Kalmar, K., Seel, R., Greenwald, B., Zafonte, R., Demarest, D., Brunner, R., & Kaelin, D. (2014). Zolpidem and restoration of consciousness. Am J Phys Med Rehabil, 93(2), 101–113. 10.1097/PHM.0000000000000069

68. Yeo, B. T., Krienen, F. M., Sepulcre, J., Sabuncu, M. R., Lashkari, D., Hollinshead, M., Roffman, J. L., Smoller, J. W., Zollei, L., Polimeni, J. R., Fischl, B., Liu, H., & Buckner, R. L. (2011). The organization of the human cerebral cortex estimated by intrinsic functional connectivity. J Neurophysiol, 106(3), 1125–1165. 10.1152/jn.00338.2011

