## Supplemental Figures for "Subcortical Hubs of Brain Networks Sustaining Human Consciousness"

### Supplemental Materials

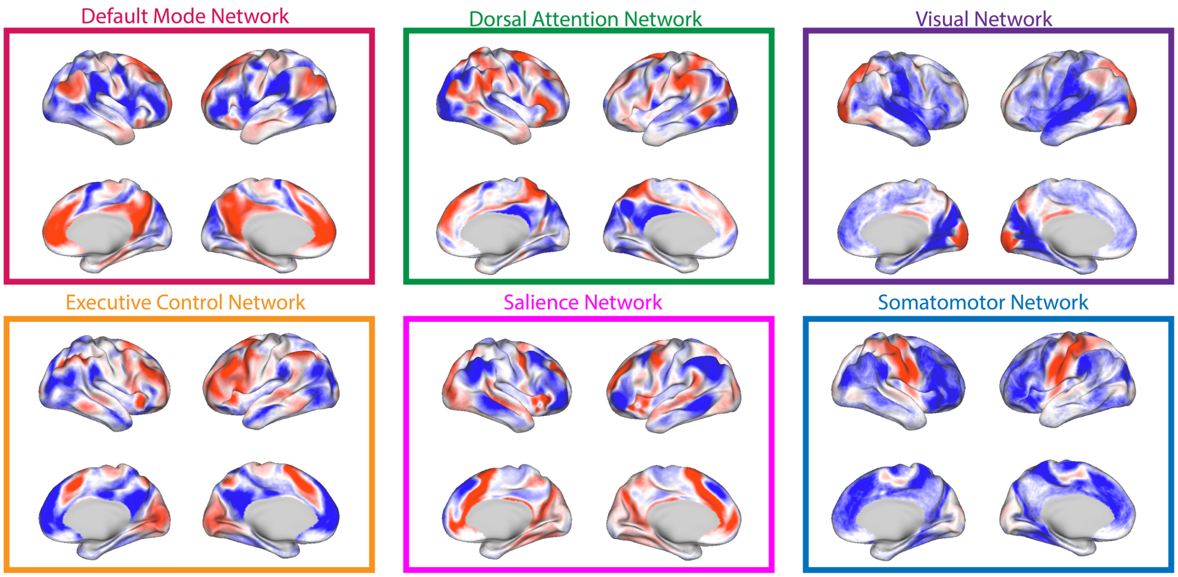


**Figure S1: Cortical maps of the six canonical networks,** namely the default mode network (DMN), the executive control network (ECN), the dorsal attention network (DAN), the salience network (SN), the visual network (VN) and the somatomotor network (SMN).

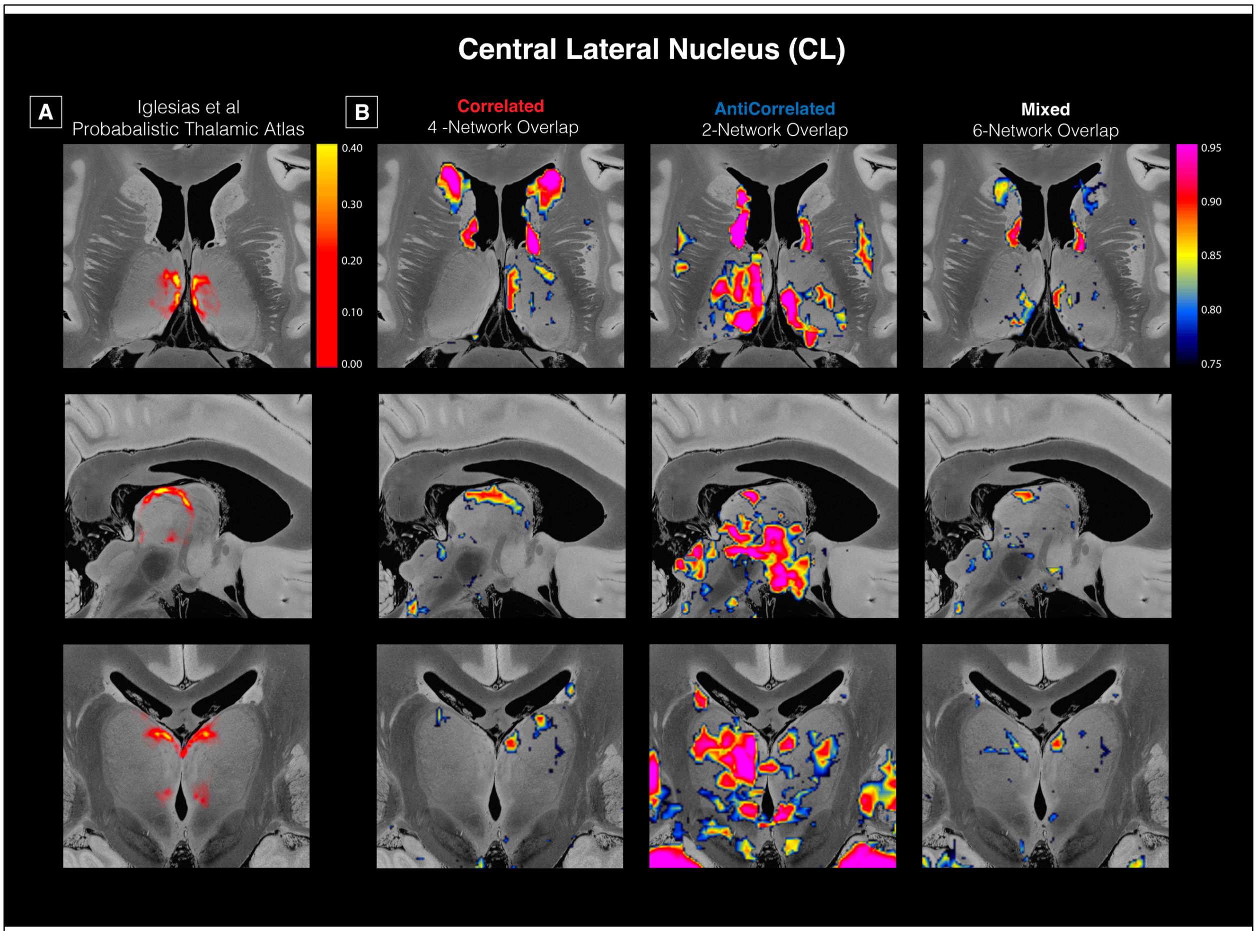


**Figure S2: Mixed connectivity findings within the central lateral thalamic nucleus.** (A) shows the right hemispheric probabilistic nuclei as defined in the thalamic segmentation atlas (Iglesias et al., 2018). (B) shows the 4-network correlated contour map (DMN, DAN, SN, VIS), the 2-network anticorrelated contour map (ECN and SMN), and the 6-network mixed contour map.

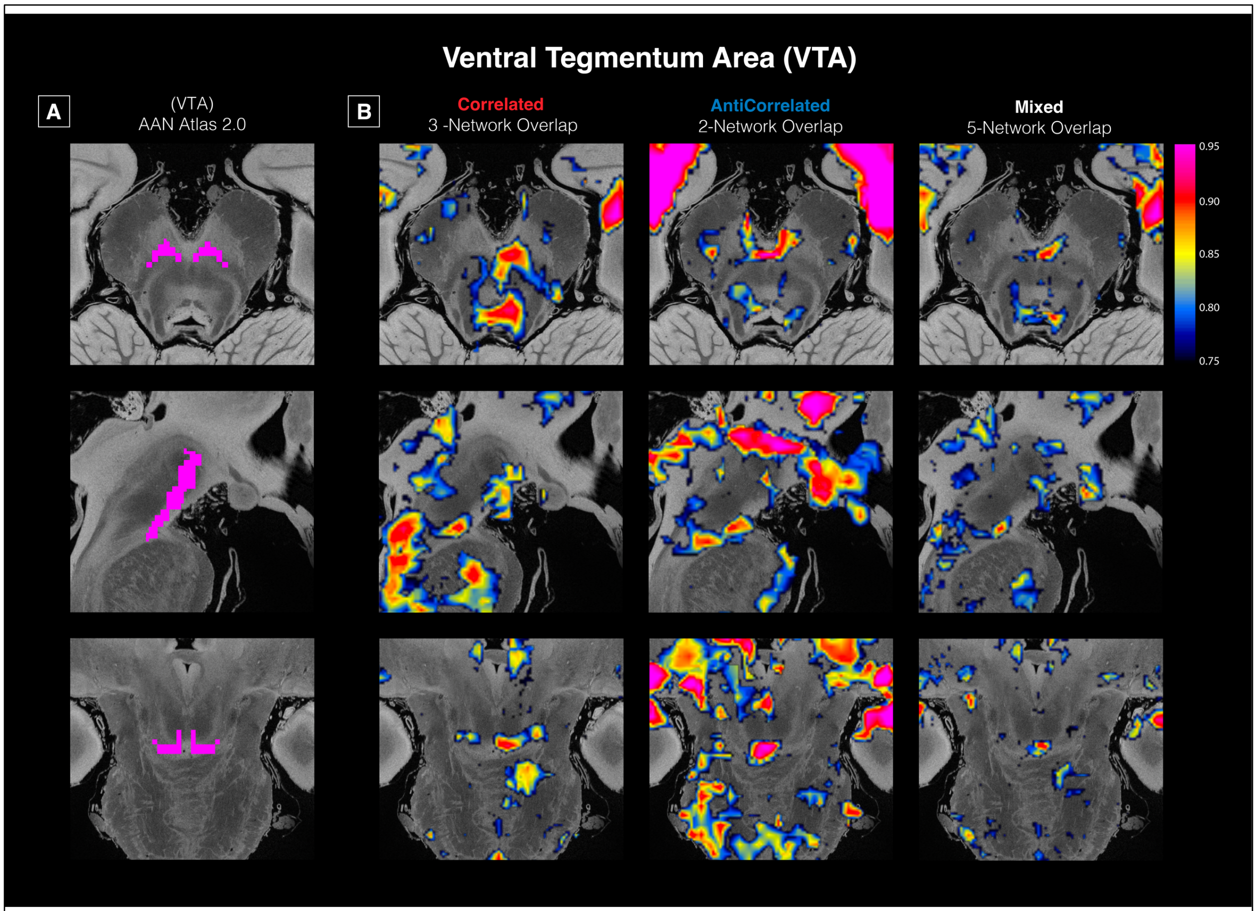


**Figure S3: Mixed connectivity findings within the VTA.** (A) shows the location of the VTA defined by the Harvard Ascending Arousal Network Atlas (Edlow, 2012). (B) shows the 3-network correlated contour map (DMN, ECN, SN), the 2-network anticorrelated contour map (DAN and SMN), and the 5-network mixed contour map (All but VN).

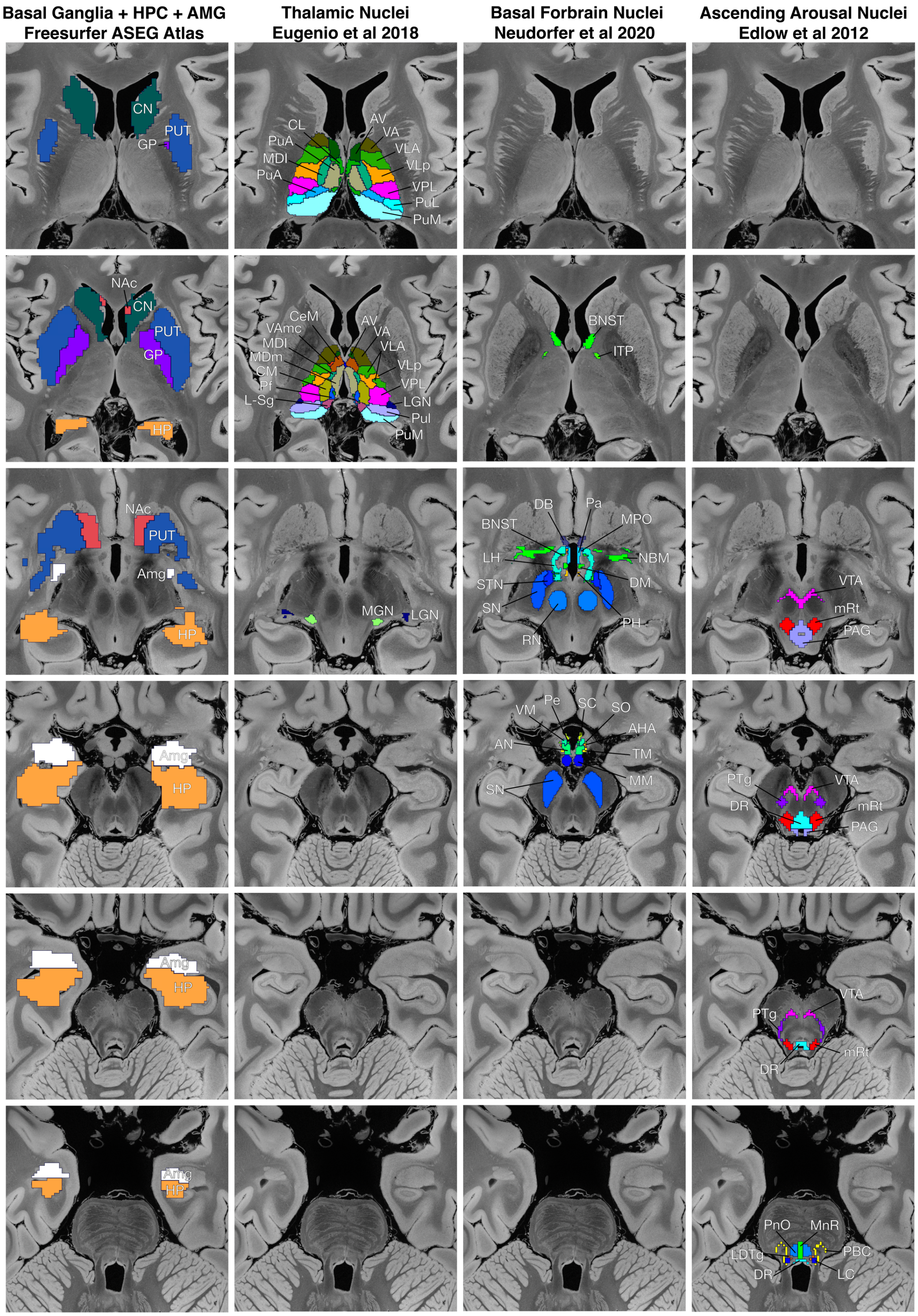


**Figure S4:** Anatomical nuclei locations for 6 axial frames within the subcortex. Each column shows the nuclei definitions for a single atlas including the ASEG Freesurfer atlas, a thalamic parcellation atlas (Eugenio et al 2018), a basal forebrain atlas (Neudorfer et al 2020), and an ascending arousal nuclei atlas (Edlow et al 2012).
